# Characterization of *Salmonella typhi* from Shallow Well Water in Sogomo, Eldoret, Kenya and its Antimicrobial Sensitivity to *Moringa oleifera* Seed Extract

**DOI:** 10.1101/2024.12.04.626862

**Authors:** Cheronoh Joy, Salinah Rono, Jelimo Emily, Luvandale Ramadhan, Kalule Francis

**Affiliations:** University of Eldoret Laboratory, Department of Biological Sciences, School of Science, University of Eldoret; Kenya Institute of Primate Research, Department of One Health and Emerging Infections; Zoetis ALPHA Plus initiative, Mercuriusstraat 20, B-1930, Zaventem, Belgium

**Author notes:** Corresponding author: Joy Cheronoh.

**Keywords:** *Salmonella typhi*, Antimicrobial Sensitivity, *Moringa oleifera*, Water contamination

## Abstract

*Salmonella typhi* is a gram negative, rod-shaped bacterium that cause typhoid fever disease, a significant cause of human mortality especially in the developing countries. This illness occurs after ingestion of *Salmonella typhi* bacteria in contaminated water or food. This study aimed to isolate and identify *Salmonella typhi* from shallow well waters and assess its sensitivity to *Moringa oleifera* seed extract. Twelve wells were sampled randomly, and samples collected. 10mls of each sample were measured and put in twelve different sterile test tubes. Dilutions were done serially by diluting 1 ml of each sample in tubes having 9mls of distilled water. This was labelled 10-1 dilution. Further dilutions were done to dilution factor of 10-3. Inoculation was done in *Salmonella Shigella* agar (SSA) and incubation done at 35 C for 24 hours. Re-inoculation was done in SSA by streaking to obtain pure isolates. These pure isolates were confirmed by biochemical tests (catalase, methyl red and Triple Sugar Iron tests). Anti-microbial sensitivity test was performed using *Moringa oleifera* seed extract and ofloxacin and norfloxacin drugs of antibiotic disc were used as positive and negative controls respectively. Poor method of human waste disposal, condition of sewage systems and inaccurate water treatment methods associate to well water contamination by *Salmonella typhi* and thus transmission. Results showed that 3 out of 12 water samples tested positive for *Salmonella typhi*. This gave a rate of 25% contamination of shallow wells in Sogomo by this bacterium. The bacterium was sensitive to *Moringa oleifera* seed extract, with sensitivity increasing with higher extract concentrations. The results of the measurements of zones of inhibition were represented using graphs. The study highlights the need for public health education among the residents of Sogomo on proper sewage management, waste disposal and water treatment to prevent contamination.

## 1.0 Introduction

*Salmonella* is a gram negative, rod shaped, non-spore forming bacterium that belong to the family of *Enterobacteriaceae*. All serovars of *Salmonellae* are capsulated except for *S. typhi* (Perilla, 2003). *S. typhi* is an example of the existing serovars that make up *Salmonella enterica. Salmonella typhi* bacterium causes typhoid fever in humans. Most human mortalities have been caused by this bacterium in the developing countries whose sanitation is poor. This bacterium has human as its reservoir and is spread through contaminated food and water via the fecal-oral route. Emergence of Multi-Drug Resistance (MDR) strains make eradication of typhoid fever almost impossible (Den *et. al*; 2007)

Human typhoid occurs after ingestion of *Salmonella typhi* bacterium usually in contaminated animal products, water or from infected individuals (Parry *et. al*; 2002, WHO, 2008). Typhoid fever shows at one to two weeks following ingestion of *Salmonella typhi*. Fever, headache, nausea, and abdominal pains are common symptoms although diarrhea can still occur, but it mostly affects the immunocompromised individuals (Parry *et. Al*, 2002. WHO, 2005). Management of *Salmonella typhi* has shown to be almost impossible due to its ability to modify its phenotypic and genotypic properties in response to environmental changes (Sanal *et. al*; 2006). It can survive partial cooking and passage through gastric acid barrier (Sanal *et. al*., 2006). This explains how it eventually enters the human intestines. *Salmonella* can thrive well in anaerobic and aerobic conditions; they have adapted to use nitrogen as the alternative source of electron acceptor instead of oxygen (Sanal *et. al*, 2006). Its isolation and identification procedures involve; selective enrichment, selective plating, and confirmatory tests (ISO, 2002). *Salmonella typhi* is acquired through contaminated water and food. Mary Mallon, a cook in Manhattan house was known as a typhoid carrier (Kenny& Kevin, 2014).

*Salmonella typhi* spread is higher in rural and urban areas with dense population due to crowding, poor sanitation, and hygiene of these areas (Hruday *et. al*., 2006). Typhoid fever estimates of the year 2000 suggested approximately 21.5 million infections and over 200,000 deaths around the globe each year (Crump *et. al*., 2004, Bhan *et. al*., 2008). It is thus considered a serious infectious disease and thus has become a threat to public health. Concerns on fast and extensive emergence of resistance to multiple antibiotics also rise (Feng, 20; Akinyemi *et. al*., 2005).

## 2. Materials and Methods

### 2.1 Study Area

The study was carried out in Sogomo area next to the University of Eldoret in Kenya. It was purposely selected due to the accessibility of the shallow wells in several residential homes of students. Sogomo is located Eldoret-Ziwa Road, besides University of Eldoret, Moiben constituency, Uasin-Gishu county. The water sources in this area are tap water, rain water and shallow wells which are commonly used. Typhoid fever has proven to be common waterborne disease in the area.

### 2.2 Study design

Experimental research design was used in the study. Water samples were collected from various wells in the study area where the sampling points were chosen using simple random method of households and residential premises. Analysis was done in University of Eldoret Botany Lab 2.

### 2.3 Sampling and data collection

Water samples (12) were collected by simple random method from six wells in Sogomo area at intervals of time; morning, noon and evening hours based on the design. This was done over a period of one day. The samples were put in sterile water bottles and labelling done appropriately, indicating the site and date of collection. The samples were then transported to the University of Eldoret microbiological laboratory, Botany lab II for analysis.

### 2.4 Media preparation and Samples Preparation

#### 2.4.1 Preparation of Salmonella -Shigella Agar

Based on the manufacturer’s instructions, Salmonella -Shigella Agar (12.604g) was dissolved in 250ml of distilled water to prepare the growth medium. The mixture was then heated to boil on a hot plate to dissolve the mixture completely. It was then allowed to cool to 50°C.

#### 2.4.2 Preparation of water samples for inoculation

10 ml from each collected samples of water were measured and put into 6 different sterile test tubes. 1 ml from each test tube was measured and diluted in a test tube containing 9 ml distilled water. This was labelled 10-1 dilution. 10-2was obtained by transferring 1 ml of the mixture to a second test tube containing 9 ml of distilled water. The 10-3 dilution was then achieved by putting 1ml of the succeeding dilutions into the tubes containing 9 ml of distilled water.

### 2.5 Culturing of the prepared samples

After the three dilutions were made, 1ml of the last dilution were transferred to the petri dishes containing Salmonella-Shigella Agar using a sterile pipette. Two replicates were made for each sample. Swirling was done and the mixture was left to cool. The petri dishes were then inverted and incubated at 35^0^C for 24 hours. After incubation, colonies were observed from the petri dishes that tested positive.

#### 2.5.1 Sub culturing

After inoculation for 24 hours, distinct colonies from primary cultures were streaked on Salmonella-Shigella Agar. Incubation was again done for 24 hours at 35°C after which observation for colonies was made. Colonies that were colourless with black centers were identified and characterized based on the colony morphology, cultural characteristics, Gram staining reaction, microscopy and confirmed by biochemical tests (Neil *et al*., Bhatta *et al*., 2005)

#### 2.5.2 Gram staining and Microscopy

A sterile wire loop was used to place a loop full of the bacterial culture on the slide and spread to make a thin smear. The dry smear was then heat-fixed by passing over a hot plate 3 times after which staining was done. After standard staining, the bacterial smear was washed and blot-dried using a blotting paper and examined under a light microscope (Olympus) at high power objective (100x) using oil immersion for morphological characteristics and staining characteristics.

#### 2.5.3 Biochemical Characterization

The colonies that were presumed to be Salmonella based on the cultural and morphological characteristics were subjected to Triple Sugar Iron test, catalase test and methyl red test. Identification of *Salmonella typhi* were based on the positive biochemical tests.

### 2.6 Antimicrobial Sensitivity Test

#### 2.6.1 Preparation of *M. oleifera* seed extract

50g of *M. oleifera* seeds were measured and grounded into fine powder. It was then soaked in 96% ethanol for three days. Filtration was then done using a filter paper to separate the filtrate from the grounded seeds residues. The recovered filtrate was taken to Chemistry lab 3 for concentration in a vacuum evaporator (figure 1), it was finally open dried on a plate for 3 days.

**Figure 1:**
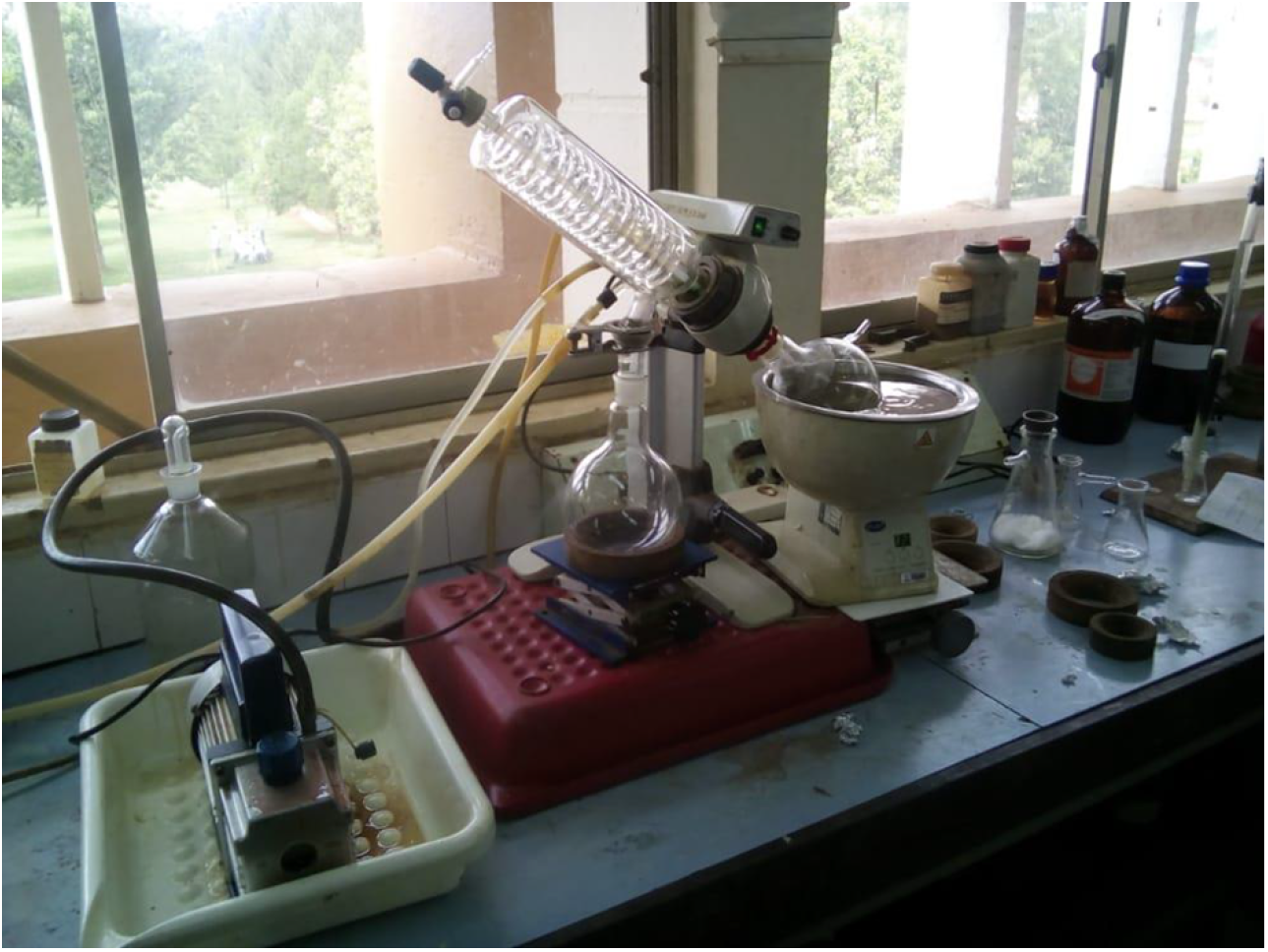
Concentration of the *M. oleifera* seed extract through a vacuum evaporator.

#### 2.6.2 Antimicrobial test procedure

Punched filter papers were dipped in the dry absolute and in diluted *M. oleifera* seed extract separately and left to stand for 1 hour. Three pieces from each concentration were then put individually with a sterile forceps on *SS*A plates containing *Salmonella* culture. Norfloxacin and Ofloxacin drugs of antibiotic disc were used as positive and negative controls respectively based on the Kirby Bauer disc diffusion method (Bauer et al., 2006). Incubation was done at 35^0^C for 24 hours. Zones of inhibition were measured in millimeters (mm). The antibiotic agents’ susceptibilities were evaluated against that of *M. oleifera* seed extract and analysis of the results done.

### 2.7 Data quality and analysis

The data collected was processed by editing, coding, and summarizing into tables in an excel spread sheet. Data was edited to check for completeness and accuracy of the reports. Graphs were used to show the differences in sensitivity of *Salmonella typhi* to *Moringa oleifera* seed extract to that of the negative and positive controls.

### 2.8 Ethical Approval

The conducted research is not related to either human or animal use.

## 3.0 Study Results

### 3.1 Isolation and morphological characterization

Isolation of *S. typhi* was done from the shallow well water. Three samples tested positive (Figure 2)

**Figure 2.**
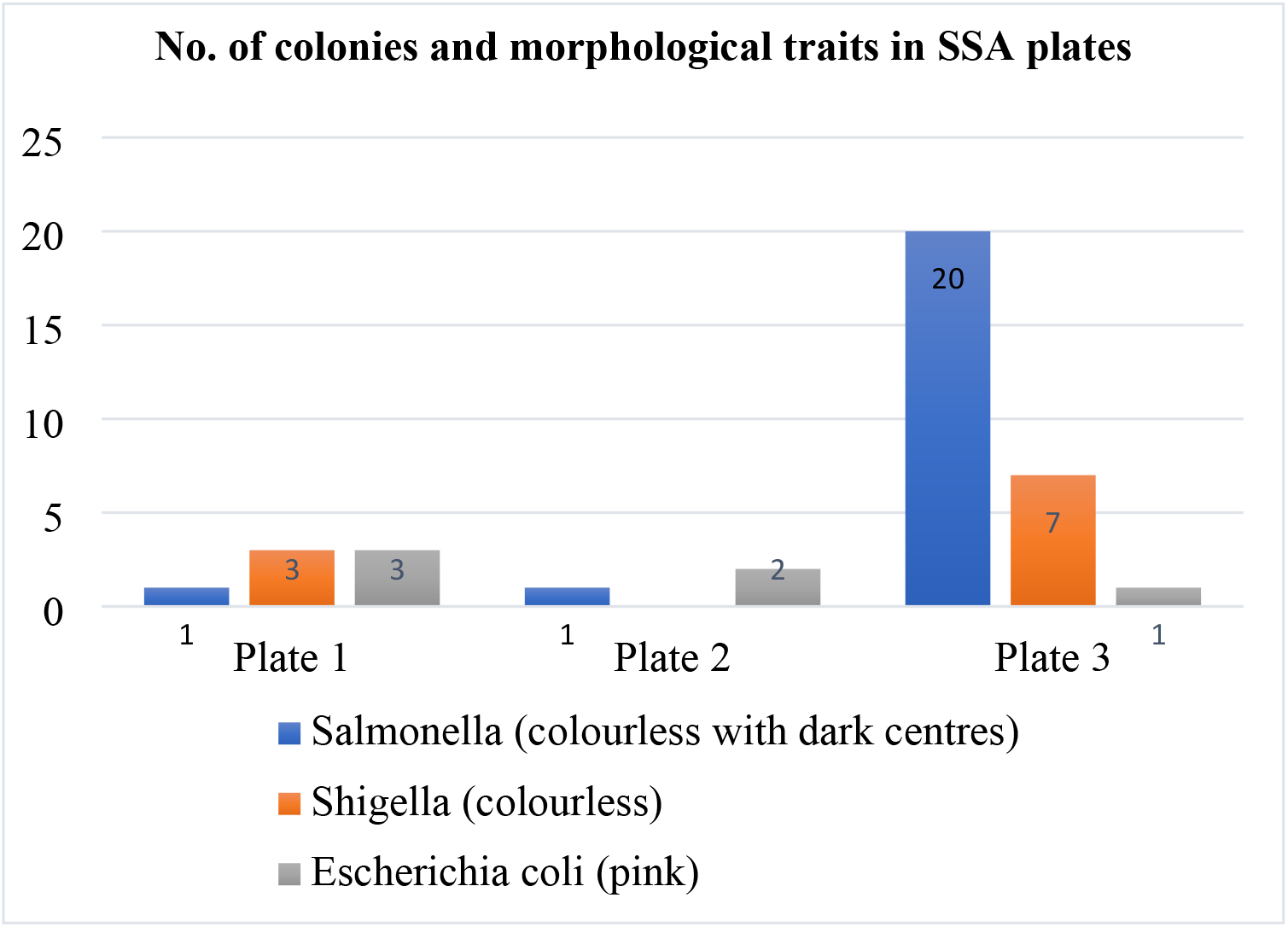
shows the total number of plates with the corresponding number of colonies in Salmonella Shigella Agar (SSA)

SSA is a differential, selective medium used for isolation of *Salmonella* and *Shigella* species from pathological specimens and suspected food. Its test sugar is a lactose, a fermentable carbohydrate. Its fermentation results in acid production and thus lactose-fermenting organisms red colonies while the non-lactose fermenters grow as colourless colonies. *Salmonella* is one of the non-lactose fermenters, that can produce hydrogen sulphide gas. It thus appears colourless with black centre. *Shigella* on another hand is incapable of both lactose fermentation and hydrogen Sulphide gas production. It thus appears colourless.

Based on the results (figure 2), there was growth of both lactose fermenter and non-lactose fermenters. Pink colonies were a characteristic of successful lactose fermentation, which presumptive *Escherichia coli* and *Klebsiella* show. The colourless colonies showed growth of *Shigella* whereas the colourless colonies with black centre showed growth of *Salmonella*.

Since *Salmonella* was the bacteria of interest, one of the colourless colonies with black centers that appeared singly, was picked, and re-inoculated into Salmonella Shigella Agar, by streaking. This was done to obtain a pure colony. The resulting pure colonies were identified, and further studied and confirmed with biochemical tests.

### 3.2 Biochemical test results

The biochemical tests that were used to achieve the confirmatory tests on *Salmonella typhi* were catalase, methyl red and the Triple Sugar Iron tests (figure 3).

**Figure 3:**
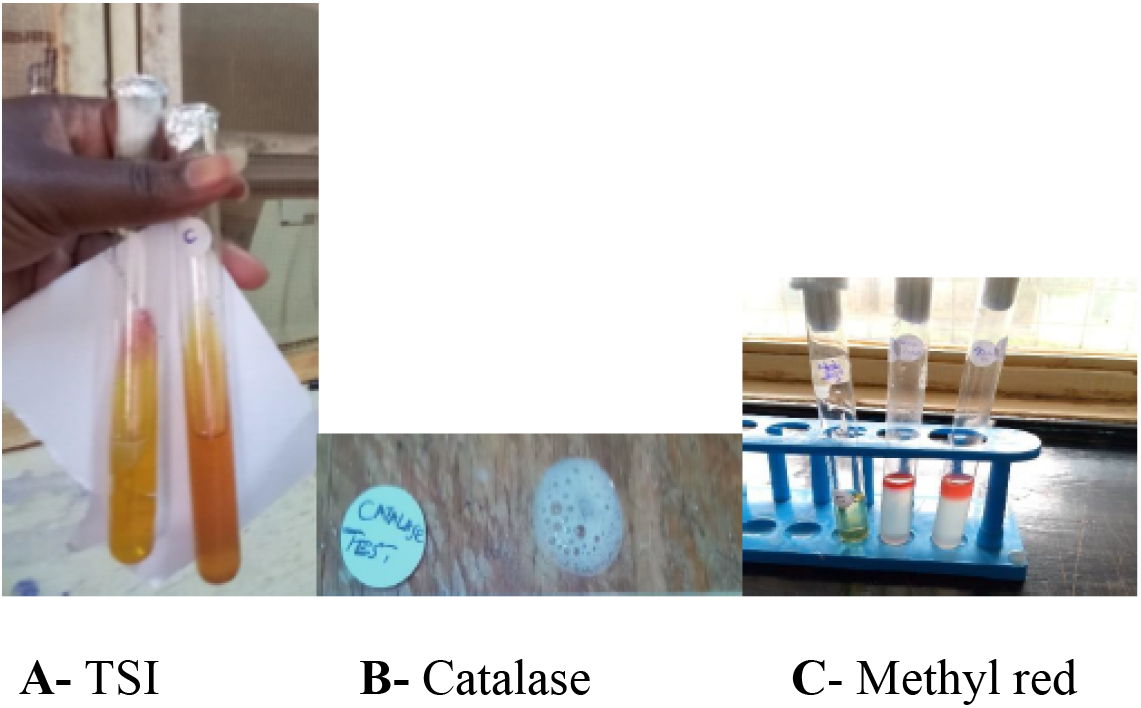
Biochemical tests (Triple Sugar Iron agar, Catalase and Methyl red) results of *Salmonella typhi*.

### 3.3. Antimicrobial Sensitivity test results

From the sensitivity test done, susceptibility and resistance of *Salmonella typhi* to each concentration of *M. oleifera* seed extract was determined. A ruler was used to measure the zones of inhibition (from the center of the treatment to the point of circumference of the zone where a distinct edge was present). The results were as shown (figure 4 and Table 1).

**Table 1:** Antimicrobial sensitivity test results for absolute and diluted *Moringa oleifera* seed extract.

*Salmonella typhi* (Zone diameter, nearest whole mm)
| Substance | Zone diameter |  |  |  |
| --- | --- | --- | --- | --- |
|  | R1 | R2 | Mean | SD |
| Absolute *M. oleifera* | 14 | 13 | 13.5 | 0.7 |
| *M. oleifera* distilled water 10<sup>5</sup> dilution | 6 | 8 | 7 | 1.4 |
| Control positive | 26 | 24 | 25 | 1.4 |
| Control negative | 2 | 5 | 3.5 | 2.1 |

**Figure 4:**
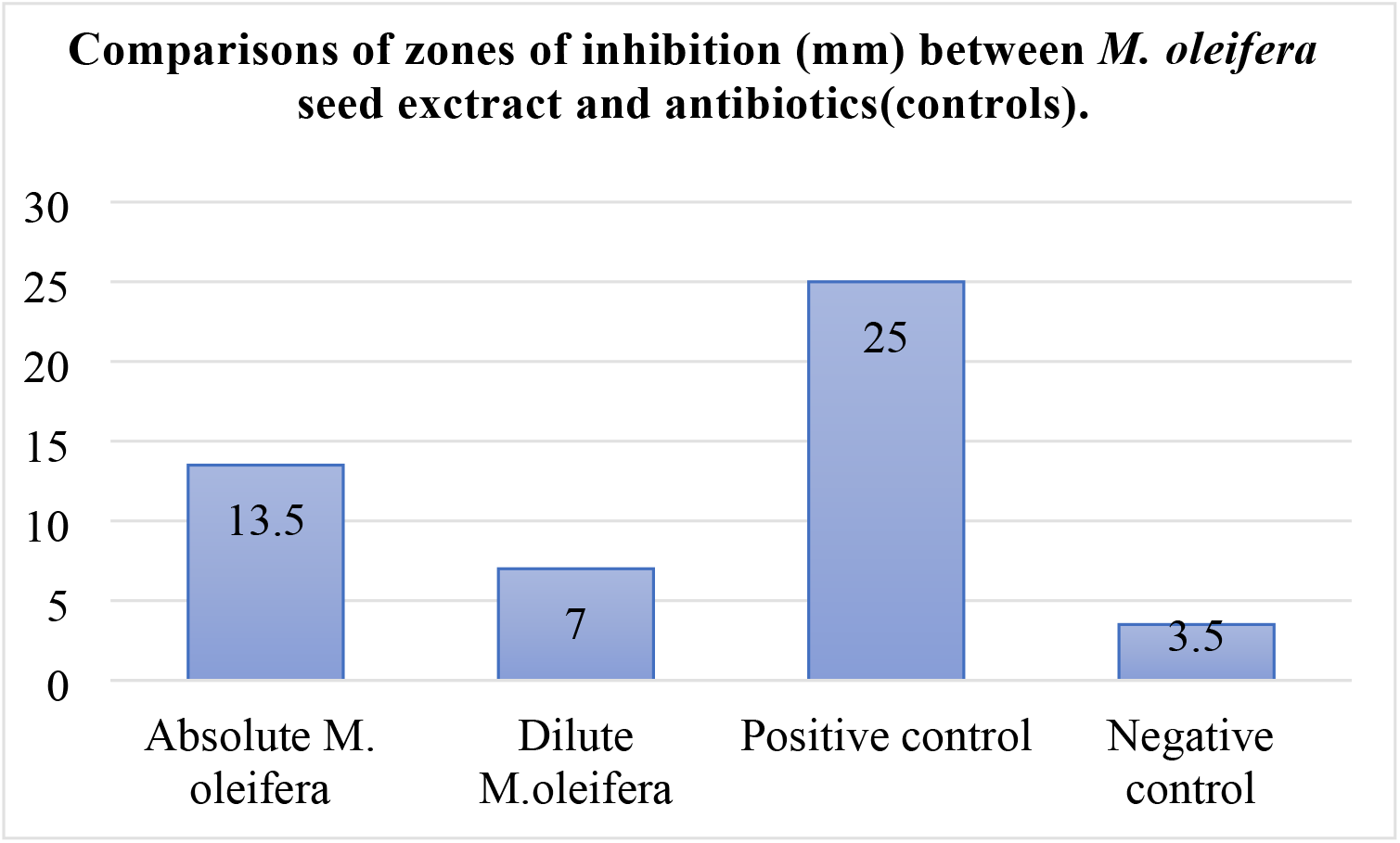
A graph showing comparison of zones of inhibition (mm) between *M. oleifera* and antibiotic controls.

The F-critical value is 19 and the P-value is 0.01 at 0.05% level of significance, for the rows that represent different *M. oleifera* seed extract concentrations and antibiotics (positive and negative controls). There is a significant difference between the zones of inhibition (supplementary information-Appendix 1)

For the columns, F-critical value is 18.5 and the P-value is 0.6 at 0.05% level of significance for the two replicates. Therefore, there is no significant difference between the zones of inhibitions.

## 4.0 Discussion

The study highlights presence of *Salmonella typhi* from shallow wells. There were three plates that tested positive for *S. typhi* out of the 12 sampled shallow wells. This gives a rate of approximately 25% of contamination of shallow wells in Sogomo area. This bacterial isolate is an enteric pathogen that is associated with different food and water-borne illnesses. The current study aligns with the previous study of Paul *et al*, 2013 that ground water are contaminated with enteric pathogens. Their records from United State and Canadian studies (1990-2013) showed existence of *Salmonella* in shallow wells. Water is a known as a common vehicle for the transmission of typhoidal *Salmonella*.

The 25% positivity rate of contamination gives sufficient evidence that water from wells in Sogomo is not safe for drinking. This agrees with the study of Olajubu & Ogonika, 2014, which reported that the presence of enteric pathogens in water often reflects faecal pollution. Surface run-off, poor disposal of human waste, inaccurate water treatment methods, including poor sewage drainage system and effluent from the septic (leak to fields can reach the shallow waters) could be the factors contributing to the existence of enteric pathogens in well water.

*Salmonella typhi* showed to be susceptible to *M. oleifera* seed ethanoic extract. This explains the bacteriostatic ability of the extract to this bacterium as seen in the growth inhibition. *Moringa oleifera* seed extract target the cell wall of *Salmonella typhi* bacterium thus killing the bacteria producing a clear zone around the punched disc embedded in the extract. A zone of inhibition of 14 mm was measured when *Salmonella typhi* was subjected to the absolute *Moringa oleifera* seed extract. Antagonistically, the aqueous form of the seed extract inhibited the growth of *Salmonella typhi* by 6mm. It is evident that zones of inhibition increase with increase in the concentration of the seed extract. The results in this current study agree with previous study of Jaganthan *et al*, 2009 that *M. oleifera* seed extract is sensitive to a wide range of enteric gram-negative bacteria including *E. coli, Shigella* and *Salmonella*. The bioactive compounds of the *Moringa oleifera* seed extract may explain the chemical reaction on *Salmonella typhi* (Oluduro, 2010)

The ethanol extract at high concentration of *M. oleifera* was efficient in inhibition of *S. typhi* than in low concentration. This is similar with the earlier results of Doughari *et al*.,2007 that, ethanoic extract of *M. oleifera* demonstrates a high activity while the aqueous extract showed the least activity against *S. typhi*.

## 5.0 Conclusion and Recommendations

This study established that, water from the shallow wells in Sogomo are contaminated with *Salmonella typhi* as shown by the plates that tested positive. 3 out of 12 samples tested positive giving a 25% rate of contamination of shallows well by *Salmonella typhi* in Sogomo area in Eldoret. The poor water treatment method, poor method of human waste disposal and the leakages of sewage systems could be the contributing factors to the presence of this bacterium in the water. Additionally, *Salmonella typhi* isolate from the shallow wells in Sogomo area is sensitive to *Moringa oleifera* seed extract. Its sensitivity increases with increase in seed extract concentration. The inhibition zones of *Moringa oleifera* seed extract are close to the inhibitory zones of control antibiotics and thus, this contributes to significant addition to the development of new chemicals that can be used to control different bacteria.

Further in-depth research on the reliability and efficacy of different extraction methods of *Moringa oleifera* seed extract should be explored, Sensitivity of gram-positive bacteria against *Moringa oleifera* seed extract should also be evaluated and there is need for use of *Moringa oleifera* seeds for well water treatment in affected areas such as Sogomo water wells to prevent further multiplication of the bacteria in wells since the seeds are cheaply available on the local markets and are safe.

## Acknowledgments

The authors are grateful to the staff of Biological Sciences department at the School of Science, University of Eldoret, and the University of Eldoret laboratory staff for the support in ensuring the success of this study.

## Funding information

The authors received no funding for this study.

## Conflict of interest

The authors state no conflict of interest.

## Data availability statement

The datasets from this study are available from the corresponding author upon reasonable request. However additional information has been provided as supplementary information.

## Notes

### Competing Interest Statement

The authors have declared no competing interest.

## References

Amen, D. (2014), Anti-microbial resistance for enteric pathogens isolated from acute gastroenteritis patients. World Journal 1, 1–14

Ang, J. Y. E., and Asmar, B. I. (2004). Antibacterial resistance. Indian Journal of Pediatrics 71, 229–239.

Ashwini, C., Nambi, P., Senthur, V., Rama Subramanian, K., Abdul Ghafur, and Thirunarayan, N. A. (2013) Antimicrobial susceptibility of Salmonella enterica serovar in a tertiary care hospital Southern India, The Journal of Medical Research, 137(4): 800–802.

Backer, H. (2002, Water disinfection for international and wilderness travellers. Clinical journal for infectious disease. 34(3). 355–364.

Bartholomew JW, Mittwer T, The Gramstain. Bacteriological views. 1952 Mar; [PubMed PMID: 14925025]

Bhutta Z. (2006) Current concepts in the diagnosis and treatment of typhoid fever, British Medical Journal 333, 78–82.

CIDRAP Center for infectious disease research and policy, academic health centre, University of minnesotta. https://www.Cidrap/contents,Fs/food/disease/cause/Salmonella

Clinical and Laboratory Standards Institute CLSI. (2007), Diagnostics and treatment of serious antimicrobial resistance of Shigella species, Infectious disease IV (4), 183–200

Crump, J., Luby, S. and Bhatnagar, S. (2005), Typhoid and paratyphoid fever. Lancet 366, 749–762.

Den, W., Shian-ren, I., Plunkett, G., Mayhew, G.F., Rose. D.J.s, Burland, V. and Voula, K. (2003). “Comparative genomics of Salmonella enterica serovar typhi strains type 2 and ct 18.” Journal of Bacteriology, volume 185, 2330–2337

Doughari. J., Micah P. & Nirmal De, (2015). Anti-bacterial activity of the Moringa oleifera on S. typhi.

Feng, Y. C. (2000). The epidemiology of typhoid fever in the Dong Thap province, Mekong. Delta region of Vietnam. The American Journal of Tropical Medicine and Hygiene, 62, 644–648

Hruday, S.E., and E.J., and Pollard. S.J. (2006), Risk Management for assuring safe drinking water. Environment International, 32(8), 984–957

ISO6579, (200), Microbiology General guidance on methods for detection of Salmonella. International Organization of Standardization, Geneve, Switzerland.

Jackson, BR., Igbal. S.N and Mahon, B. (2015). “Updated recommendation for the use of typhoid vaccine advisory committee on immunization practices, United States.” Morbidity and mortality weekly report, 64(11): 305–308

Kenny and Kevin (2014). The American Irish. A History Routledge. P. 187

Libenson L, Mcilroy AP, On the mechanism of gram stain. The Journal of infectious diseases. 1955 Jul-Aug

Midala, T., Agina. S.A., and Maselle, S. Y., (2010). Bacteria isolated from bloodstream infectious at a tertiary hospital in Dares Salaam, Tanzania antimicrobial resistance of isolates. South African Medical Journal 100(12), 835–8

Morpeth, S. C, Ramadhani, H. O. and Crump, J. A (2009), Invasive non-typhi Salmonella disease in Africa. Clinical infectious diseases 49, 606–611

O. A. Oluduro, T.O. Idowu, B.I. Aderiye, O. Fameruwa and O.O. Omoboye. Evaluation of Antimicrobial Potential of Crude Extract OF Moringa oleifera seed on Orthopaedic Wound Isolates and Characterization of Phenylmethanamine and Benzyl Isothiocyanate Derivatives.

O’ Toole GA, Classic Spotlight: How the Gram Stain Works. Journal of bacteriology. 2016 Dec 1.

Paul D. Hynds, M. K. Thomas, Katarina D. M. Contamination of groundwater systems in the US and Canada by Enteric Pathogens, 1990–2013

Parry, C. M., Hien, T. T., Dougan, G., Whiter, N. J., and Farvar J. J. (2002). Typhoid fever, National English Medical Journal 347, 1770–1782

Sanal, O., Turul, T., De boer, T., Van, D.V., Yalan, I and Tezlan, I (2006), presentation of interleukin-12/-23 receptor beta 1 deficiency with various clinical symptoms of Salmonella infections. Journal of Clinical Immunology 26, 1–4

Sources of Fecal Bacterial Pollution of Environmental Waters. JPS Cabral, 2010

WHO. (2002), Water sanitation and health report. Water and Sanitation Programme. WHO/WHS/WWD/TA8.

Winfred, W., and Julia M. (2008), Comprehensive Geography. Longhorn Publishers Limited Nairobi, Kenya.

